# Quantitative analysis of non-histone lysine methylation sites and lysine demethylases in breast cancer cell lines

**DOI:** 10.1101/2024.09.18.613658

**Authors:** Christine A. Berryhill, Taylor N. Evans, Emma H. Doud, Whitney R. Smith-Kinnaman, Jocelyne N. Hanquier, Amber L. Mosley, Evan M. Cornett

## Abstract

Growing evidence shows that lysine methylation is a widespread protein post-translational modification that regulates protein function on histone and non-histone proteins. Numerous studies have demonstrated that dysregulation of lysine methylation mediators contributes to cancer growth and chemotherapeutic resistance. While changes in histone methylation are well documented with extensive analytical techniques available, there is a lack of high-throughput methods to reproducibly quantify changes in the abundances of the mediators of lysine methylation and non-histone lysine methylation (Kme) simultaneously across multiple samples. Recent studies by our group and others have demonstrated that antibody enrichment is not required to detect lysine methylation, prompting us to investigate the use of Tandem Mass Tag (TMT) labeling for global Kme quantification sans antibody enrichment in four different breast cancer cell lines (MCF-7, MDA-MB-231, HCC1806, and MCF10A). To improve the quantification of KDMs, we incorporated a lysine demethylase (KDM) isobaric trigger channel, which enabled 96% of all KDMs to be quantified while simultaneously quantifying 326 Kme sites. Overall, 142 differentially abundant Kme sites and eight differentially abundant KDMs were identified between the four cell lines, revealing cell line-specific patterning.

## INTRODUCTION

Lysine methylation (Kme) is a common post-translational modification (PTM) that occurs when one to three methyl groups are added to the ε- amine of a lysine sidechain by lysine methyltransferases (KMTs). Kme is a reversible PTM, and its removal is mediated by lysine demethylases (KDMs). Proteins with a characterized methyllysine recognition domain (readers) interact with Kme sites and perform downstream molecular functions. The role of Kme has been predominantly studied as an epigenetic regulator in the context of histone proteins but work over the past two decades has uncovered over 10,000 Kme sites on close to 5,000 proteins in the human proteome^1^. Characterized non-histone Kme sites regulate protein-protein interactions, subcellular localization, DNA-protein interactions, RNA-protein interactions, activity, and stability^2^.

In addition to recent developments characterizing the molecular function of Kme, many studies have shown a clear connection between Kme site dysregulation and human diseases, including breast cancer^3–6^. Breast cancer is the second leading cause of cancer death in women^7^. The amplification or loss of KMTs and KDMs has been shown to contribute to breast cancer growth and drug resistance^8–10^. Considerable efforts to target KMTs and KDMs have shown promise in preclinical studies. For example, inhibitors of the KMTs SUV39H2 and EZH2 were shown to reduce the proliferation of breast cancer cells^11–13^. Modulation of Kme on histone proteins is typically the objective in these studies, but substantial evidence indicates that Kme sites on non-histone proteins play a significant role in breast cancer progression and metastasis. For example, the KMT SMYD2 regulates breast cancer metastasis by methylating BCAR3 and modulating lamellipodia^14^. Another group demonstrated that breast cancer metastasis is controlled by KMT5A-mediated monomethylation of SNIP1^15^. Despite these findings, no systematic efforts have been made to quantify the changes in non-histone lysine methylation in breast cancer cells.

The ability to reproducibly identify and quantify Kme sites in a high-throughput manner across entire proteomes remains a significant challenge, hindering our understanding of the functional consequences of specific Kme sites. Most efforts have relied on affinity reagents to enrich methylated proteins or methylated peptides^16–20^. Relative quantitation has been accomplished by coupling enrichment with isobaric labeling using stable isotope labeling by amino acids (SILAC) or tandem-mass tags (TMT)^20–22^. We recently compared several sample preparation strategies for globally profiling lysine methylation and found that enrichment of methylated proteins or peptides by affinity reagents is not required^1^. Bypassing the enrichment step reduces the sample material requirements and enables the characterization of the levels of the enzymes responsible for regulating Kme sites in the same experiment.

Connecting KMTs and KDMs to specific lysine methylation sites remains challenging. Many groups have found success by manipulating the expression of KMTs and KDMs in cells or lysates followed by enrichment of methylated proteins or peptides and tandem MS analysis. For example, SMYD2 substrates were discovered by enriching them with pan-methyllysine antibodies in differentially SILAC-labeled cell lines with upregulated or downregulated SMYD2 expression^20^. Using an alternative approach, SILAC-labeled lysates were used as substrates for SMYD2 reactions, followed by enrichment with a pan methyllysine reader protein (3xMBT)^14^. Quantifying KMTs and KDMs simultaneously with lysine methylation sites would capture any network effects perturbations have on the entire methylation signaling network. Recently, isobaric trigger channels have been employed to boost the number of defined peptides in isobaric labeling-based quantitative MS approaches, allowing confident identification and quantification of targeted peptides or proteins in an untargeted proteomics experiment^23–27^. In this proof-of-principle study, we build upon our finding that enrichment is not required for deep lysine methylome coverage and demonstrate that tandem mass tag (TMT) labeling combined with a KDM isobaric trigger channel enables relative quantification of lysine methylation sites and lysine demethylases (KDMs) simultaneously in various breast cancer cell lines.

## MATERIALS AND METHODS

### Cell Culture and Transfections

MDA-MB-231 and HCC1806 cell lines were cultured in Dulbecco’s Modified Eagle Media (DMEM)(Corning) supplemented with 10% (v/v) fetal bovine serum (Sigma) and 1x Penicillin/Streptomycin (Corning). MCF7 and HEK293T cells were cultured in RPMI media supplemented with 10% (v/v) fetal bovine serum and 1x Penicillin/Streptomycin. MCF10A was cultured in DMEM F-12 SILAC media with 5% Horse Serum, 20 ng/µL EGF (Peprotech), 0.5 mg/mL Hydrocortisone (Sigma #H-0888), 100 ng/μL Cholera Toxin, 10 µg/mL Insulin, and 1x Pen/Strep. All cells were incubated at 37 °C with 5% CO_2_. For GFP-KDM transfections,10 µg of an individual GFP-KDM with 30 µL of PEI were transfected into a 10 cm HEK293T dish. Cells were collected after 24 hrs.

### Western Blotting

Cells were grown until 90% confluent and resuspended in lysis buffer (10 mM Pipes pH 7, 300 mM sucrose, 100 mM NaCl, 3 mM MgCl2, 0.1% Triton X-100, protease inhibitor, universal nuclease (Pierce)). Lysates were run on a 6% SDS-PAGE and probed with anti-GFP (ProteinTech Catalog #50430-2-AP, 1:1000 dilution) or anti-Tubulin (Proteintech #66240-1-Ig, 1:20,000).

### Immunoprecipitation of the GFP-KDMs

Cell lysates were normalized according to the western blot results. Lysate was combined with 10 μL of GFP-Trap Agarose Magnetic beads with a final volume of 250 μL of IP Buffer (10 mM Tris pH 7.5, 150 mM NaCl, 0.5M EDTA) and incubated for 1 hr at 4°C with rotation. Beads were washed 5 times for 5 minutes with the IP Dilution buffer. Before the last wash, the beads were combined for the last wash.

### IP LC-MS/MS Sample Preparation

Beads were covered with 30 µL 8 M urea in 100 mM Tris, pH 8.5 with 5 mM tris (2-carboxyethyl)phosphine hydrochloride (Sigma-Aldrich Cat No: C4706) for 30 min at 35°C. The beads were then treated with 10 mM chloroacetamide (final concentration, Sigma Aldrich Cat No: C0267) for 30 min at room temperature in the dark. Samples were diluted with 50 mM Tris pH 8.5 (Sigma-Aldrich Cat No: 10812846001) to a final urea concentration of 2 M for overnight Trypsin/Lys-C digestion at 35°C (1 µg protease used, Mass Spectrometry grade, Promega Corporation, Cat No: V5072). After acidification, the sample was desalted by Sep-Pak (50 mg, Waters™ Cat No: WAT054955) with a wash of 1 mL 0.1% trifluoroacetic acid (TFA) followed by elution in 0.6 mL of 70% acetonitrile 0.1% formic acid (FA). Peptides were dried by speed vacuum and resuspended in 100 µL 0.1% FA for initial IP-MS analysis.

### IP LC-MS/MS and data analysis

Mass spectrometry was performed utilizing an EASY-nLC 1200 HPLC system (SCR: 014993, Thermo Fisher Scientific) coupled to Exploris 480™ mass spectrometer with FAIMSpro interface (High Field Asymmetric Ion Mobility, Thermo Fisher Scientific). 1/20^th^ of the IP was loaded onto a 25 cm EasySpray column (ES902, Thermo Fisher Scientific) at 350 nL/min. The gradient was held at 5% B for 5 minutes (Mobile phases A: 0.1% formic acid (FA), water; B: 0.1% FA, 80% Acetonitrile (Thermo Fisher Scientific Cat No: LS122500)), then increased from 4-30%B over 98 minutes; 30-80% B over 10 mins; held at 80% for 2 minutes; and dropping from 80-4% B over the final 5 min. The mass spectrometer was operated in positive ion mode, default charge state of 2, advanced peak determination on, and lock mass of 445.12003. Three FAIMS CVs were utilized (−40 CV; −55 CV; −70CV) each with a cycle time of 1.3 s and with identical MS and MS2 parameters. Precursor scans (m/z 375-1500) were done with an Orbitrap resolution of 120000, radio frequency lens% 40, automatic maximum inject time, standard automatic gain control (AGC) target, minimum MS2 intensity threshold of 5e3, monoisotopic precursor selection (MIPS) mode to peptide, including charges of 2 to 7 for fragmentation with 30 sec dynamic exclusion. MS2 scans were performed with a quadrupole isolation window of 1.6 m/z, normalized HCD of 30, 15000 Orbitrap resolution, standard AGC target, automatic maximum injection time (IT), fixed first mass of 110 m/z.

Data were analyzed in Proteome Discoverer 2.5 using a reviewed *Homo sapiens* proteome UniProt FASTA (downloaded 05/13/22; 20,292 sequences) plus common contaminants (71 sequences). SEQUEST HT searches were conducted with a maximum number of 4 missed cleavages; precursor mass tolerance of 10 ppm; and a fragment mass tolerance of 0.02 Da. The maximum number of modifications per peptide was set to 3. Dynamic modifications used for the search were 1) carbamidomethylation on cysteine (C) residues, 2) oxidation of methionine, 3) acetylation, 4) methionine loss, or 5) acetylation with methionine loss on protein N-termini. Percolator False Discovery Rate (FDR) was set to a strict setting of 0.01 and a relaxed setting of 0.05.

### Quantitative Proteomics Sample Preparation and Labeling

Cell pellets (*n*=4, total of 16 samples) were resuspended in 400 µL 8 M urea, in 100 mM Tris, pH 8.5. Samples were then sonicated on a Bioruptor® sonication system (Diagenode Inc. United States, North America Cat No: B01020001) with 30 s/30 s on/off cycles for 20 min in a water bath at 4°C. Samples were treated with 2 units of MilliporeSigma Benzonase 1016540001) and then sonication was repeated. After subsequent centrifugation at 12,000 rcf for 30 min, protein concentrations were determined by Bradford protein assay (BioRad Cat No: 5000006). 80 µg equivalent of protein from each sample were then treated with 5 mM tris (2-carboxyethyl)phosphine hydrochloride (Sigma-Aldrich Cat No: C4706) to reduce disulfide bonds for 30 min at 35 °C, then resulting free cysteine thiols were alkylated with 10 mM chloroacetamide (final concentration, Sigma Aldrich Cat No: C0267) for 30 min at room temperature in the dark. Samples were diluted with 50 mM Tris pH 8.5 (Sigma-Aldrich Cat No: 10812846001) to a final urea concentration of 2 M for overnight Trypsin/Lys-C digestion at 35°C (1:50 protease:substrate ratio, Mass Spectrometry grade, Promega Corporation, Cat No: V5072).

Digestion was halted by addition of 0.5% final v/v trifluoroacetic acid (TFA), and peptides were desalted on Waters Sep-Pak® Vac cartridges (50 mg, Waters™ Cat No: WAT054955) with a wash of 1 mL 0.1% TFA followed by elution in 0.6 mL of 70% acetonitrile 0.1% formic acid (FA). Peptides were dried by speed vacuum and resuspended in 25 µL 100 mM triethylammonium bicarbonate, pH 8.5. Each sample was then labeled for 2 hours at room temperature with 0.5 mg Tandem Mass Tag Pro (TMTpro) reagent (TMTpro Cat No: 44520, Lot no XK347989). Samples were checked to ensure labeling efficiency of >90 % and then quenched with 0.3% hydroxylamine (final v/v) at room temperature for 15 min. Labeled peptides were then mixed and dried by speed vacuum.

Approximately 1/3^rd^ of the labeled peptide mixture was fractionated using the TMT fractionation protocol of Pierce high pH basic reversed-phase peptide fractionation kit (Thermo Fisher Scientific™ Cat no 84858; with a wash of 5% acetonitrile, 0.1% triethylamine (TEA) followed by elution in 12.5%, 15%, 17.5%, 20%, 22.5%, 25%, 30%, and 70% acetonitrile, all with 0.1% triethylamine (TEA)).

### Quantitative Proteomics LC-MS/MS

1/4^th^ of each global peptide fraction was injected using an EasyNano LC1200 coupled with 25cm Aurora column (Ionopticks AUR3-25075C18-TS) on an Exploris 480 Orbitrap mass spectrometer (Thermo Fisher Scientific) with FAIMSpro installed. Peptides were eluted over a 180-minute method: Solvent B was increased from 8%-38% over 160 min, to 90% B over 10 min, held at 90% B for 5 min and decreased 8% B over 5 min (Solvent A: water, 0.1% formic acid; Solvent B: 80% acetonitrile, 0.1% formic acid). The mass spectrometer was operated in positive ion mode, advanced peak determination on, and a user-defined lock mass of 445.12003, with 3 FAIMS CVs (−45, −55, −65). A cycle time of 2 s was used for each CV and RF lens was set to 40%. MS1 parameters for each cycle were: Orbitrap resolution of 60,000, scan range of 375-1500 m/z, normalized AGC target of 300%, 50 ms max IT, minimum intensity of 5 × 10^−4^, charge state 2-8, 60 sec dynamic exclusion and excluding isotopes from fragmentation. MS2 settings were isolation of 0.7 m/z, fixed HCD of 32, Orbitrap resolution of 45,000, fixed first mass of 100, AGC target of 200%, and max IT of 120 ms.

### Quantitative Proteomics Database Searching

Data were analyzed in Proteome Discoverer 2.5 using a reviewed *Homo sapiens* proteome UniProt FASTA (downloaded 05/13/22; 20,292 sequences) plus common contaminants (71 sequences). SEQUEST HT searches were conducted with a maximum number of 5 missed cleavages; precursor mass tolerance of 10 ppm; and a fragment mass tolerance of 0.02 Da. The maximum number of modifications per peptide was set to 5. Static modifications used for the search were 1) carbamidomethylation on cysteine (C) residues. Dynamic modifications used for the search were 1) oxidation of methionine, 2) deamidation of arginine and asparagine, 3) methylation on lysine or arginine, 4) dimethylation on lysine, 5) trimethylation on lysine, 6) TMT label on lysine. Dynamic peptide and protein terminus modifications included 1) TMT label and 2) acetylation on the N-termini of peptides, 3) methionine loss or 4) acetylation with methionine loss on protein N-termini. Percolator False Discovery Rate was set to a strict setting of 0.01 and a relaxed setting of 0.05. IMP-ptm-RS node was used for all modification site localization scores. Values from both unique and razor peptides were used for quantification. In the consensus workflows, peptides were normalized by total peptide amount with no scaling. Quantification methods utilized isotopic impurity levels available from Thermo Fisher Scientific (lot numbers above). Reporter ion quantification was allowed with signal-to-noise threshold of 6 and co-isolation threshold of 30%. Resulting grouped abundance values for each sample type, abundance ratio (AR) values; and respective p-values (individual protein, ANOVA) from Proteome Discover™ were exported to Microsoft Excel for downstream pathway analysis.

### KDM Trigger Channel Proteomics LC-MS/MS

Half of the total IP digest was labeled with TMTpro label 134C as described for cell lines above. An estimated 2 µg of labeled IP peptides were mixed with one of the 1/3^rd^ unfractionated TMTpro mix from the initial cell line study (approximately 1:50). This sample was then high pH basic fractionated into 8 fractions (as described above) and each fraction was run on the Exploris 480 mass spectrometer with parameters as described above. A technical replicate was also run with the same instrument settings, but FAIMS CVs adjusted to −50, −60 and −70 V. Data analysis of the 17-plex containing trigger channel was performed as described above. Data were examined with and without the 134C channel used in the analysis.

### Data Analysis

All downstream analysis was done using R v.4.3.1. Proteins were filtered for a high FDR with more than one unique peptide, and a pseudo count of 0.01 was added to the normalized abundances to mitigate data analysis run errors. Lysine methylated peptides were filtered for a Site Localization Score (ptmRS module) greater than 0.9 and unambiguous. The Kme peptide normalized abundance values were normalized to the protein normalized abundance values for each replicate. Differentially abundance proteins and normalized Kme sites were identified using an ANOVA (*p* < 0.05) followed by Tukey pair-wise comparisons (*p* <0.05). Fold change cut-offs of |1.25| were applied. PhosphoSitePlus© data was accessed in May 2024^32^. The following packages were used for the indicated analysis: *clusterprofiler* (GO term analysis)^46^, *ggplot2* (bargraphs), *ggpubr* (stats on bargraphs), *pheatmaps* (heatmaps), *VennDiagram* (venn diagrams). The *WGCNA* R package was used for co-expression network analysis using a signed network approach^47^. A soft-threshold power of 20 (for protein WGCNA) or 16 (for Kme sites) was used. Modules with a minimum size of 30 (protein) or 20 (Kme sites) were identified using a hierarchical cluster. The output of data anlaysis is available in **Table S3**.

## RESULTS

### Generating a lysine demethylase isobaric trigger channel

Previous studies have shown that incorporating purified target proteins into an isobaric trigger channel enhances peptide levels beyond the MS1 threshold needed for selection using data-dependent acquisition (DDA) with little impact on the overall proteome coverage or quantitation^23–27^. Given the low abundance of lysine demethylases, we sought to create an isobaric trigger channel as a proof-of-concept to boost the detection and quantification of KDMs while still characterizing global lysine methylation (**Fig. 1A**). To generate the trigger channel, all 27 human KDMs were exogenously expressed as N-terminal GFP fusions in HEK293T cells. Immunoblot analysis was used to confirm expression and normalize the amount of lysate used in immunoprecipitation experiments (**Fig. 1B**). KDM5B was the only KDM without a visible band at the expected size. Individual KDMs were immunoprecipitated using magnetic GFP-Trap beads, and then all 27 KDM immunoprecipitations (IPs) were combined before on-bead trypsin digestion. The GFP-KDM IP was analyzed by LC-MS/MS, resulting in the detection of all KDMs except KDM5B (*n* = 26/27) (**Fig. 1C**).

**Figure 1:**
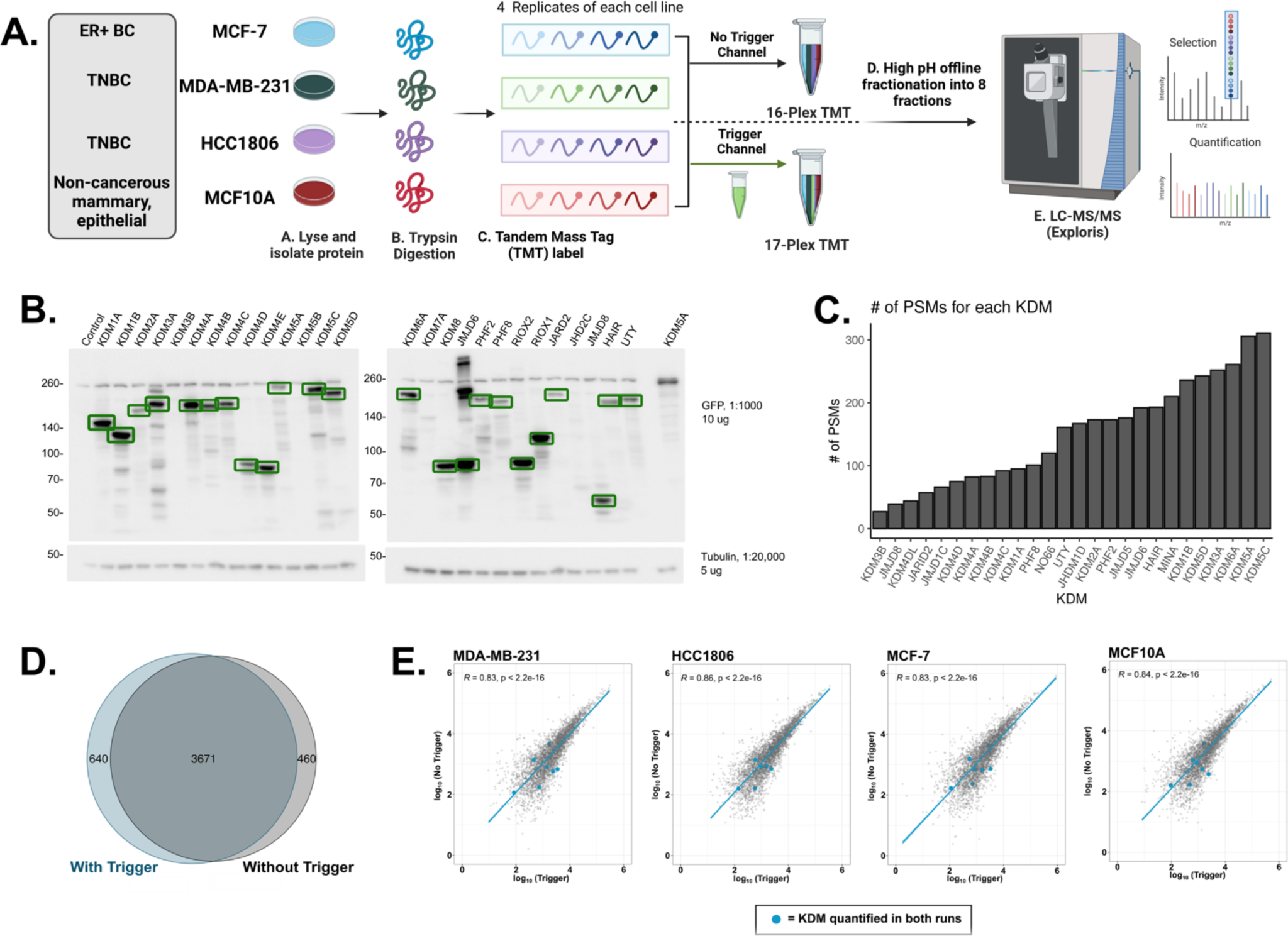
Comparison of the breast cancer proteomes with and without a lysine demethylase trigger channel. **A.** Breast cancer cell lines (MCF-7, MDA-MB-231, HCC1806, or MCF10A; *n* = 4) were collected, lysed, and trypsin/Lys-C digested. Samples were then multiplexed using tandem mass tag (TMT) labels (16-plex). The trigger channel also received its own TMT label, which was then added in a 1:50 ratio to part of the 16-plex, resulting in one 16-plex and one 17-plex. Multiplexes were subjected to high pH offline fractionation and run on the LC-MS/MS. Proteome Discoverer (2.5) was used for database searching. **B.** Western Blot of the exogenous expression of all GFP-tagged KDMs. Green boxes indicate bands at the predicted molecular weight. **C**. Bar graph depicting the number of PSMs detected for each KDM detected in the IP-MS experiment. **D.** Venn Diagram of the quantified protein groups identified with and without the trigger channel. **E.** Pearson correlation of the log_10_ average protein abundances observed with (x-axis) and without (y-axis) the trigger channel. Plots are shown by cell line and KDMs quantified in both experiments are shown as blue points.

### Adding the lysine demethylase trigger channel does not impact protein quantification

Given the successful detection of KDMs, we then inquired if the addition of the KDM trigger channel could boost the detection of KDMs without impacting the relative quantification of the whole proteome. We collected lysates (*n* = 4) from two basal-like triple-negative breast cancer (TNBC) cell lines (HCC1806 and MDA-MB-231), a Luminal A breast cancer cell line (MCF-7), and a non-cancerous, epithelial, mammary cell line (MCF10A). Each biological replicate received a tandem mass tag (TMT) label, resulting in a 16-plex sample. To test the efficacy of the KDM trigger channel, two µg of the KDM IP was TMT labeled and added to 100 μg of the 16-plex, resulting in a 17-plex sample. This ratio has been used by others and is low enough of a ratio to trigger detection but not enough to impact protein quantification. The non-trigger and KDM trigger channel experiments were fractionated offline before tandem mass spectrometry analysis (**Fig 1A**). Without the trigger channel, 142,475 peptide spectral matches (PSMs), 54,727 peptides, 4,200 detected proteins, and 4,131 quantified proteins were identified. A similar number of PSMs (126,891), peptides (47,699), detected proteins (4,384), and quantified proteins (4,311) were identified with the addition of the trigger channel (**Table S1**). Most quantified proteins (85-88%, *n* = 3,671) were identified in both experiments (**Fig 1D**).

We then compared the protein abundances between the two experiments to ensure that the inclusion of the trigger channel had no impact on the relative quantification levels of proteins. The protein abundance of the biological replicates in both experiments had a Pearson correlation coefficient equal to or greater than 0.8, demonstrating a high degree of reproducibility within each experiment (*p*< 0.05) (**Fig S1A & B**). Pearson correlation analysis of the averaged protein abundances from each cell line revealed a high degree of reproducibility between the two experiments (*n* = 3,671; R > 0.8, *p* < 0.05) (**Fig 1E**). Principal component analysis (PCA) of the protein abundances showed similar distinct clusters of the replicates for each cell line, regardless of the inclusion of the trigger channel. Notably, the three mammary epithelial cell lines (MDA-MB-231, HCC1806, and MCF10A) remained segregated from the ER+ cell line (MCF-7) (**Fig S1C & D**). Taken together, these analyses suggest the addition of the KDM trigger channel had little impact on the quantitation of the breast cancer proteomes.

### WGCNA uncovered significant differences in protein expression profiles between cell lines

We reasoned that the three distinct cell subtypes (non-cancerous, basal-like TNBC, and ER+ breast cancer) used in the MS experiments would have differentially regulated proteomes. We conducted a weighted gene correlation network analysis (WGCNA) on the differentially abundant proteins to identify clusters and characterize sub-type differences. Since the two MS experiments (with and without the trigger channel) had similar relative abundances for the proteins in common, the following proteome analysis was only conducted using the trigger channel dataset. Across the four samples, 2,290 proteins were differentially abundant (ANOVA, *p* < 0.05). In total, seventeen modules were identified (**Fig S2A & B**). Interestingly, the TNBC cell lines were markedly distinct and clustered the furthest from each other (**Fig S2C**). Furthermore, HCC1806 and MCF10A were clustered within their own branch. Previous proteomics studies have shown that HCC1806 and MCF10A cluster together, while MDA-MB-231 falls within a different cluster^28^.

Gene Ontology (GO) enrichment analysis was then performed on the most significant modules for each cell line (**Fig S2D**). Proteins associated with HCC1806 (brown module) were enriched for components of the extracellular matrixes and various fatty acid metabolic processes, including the amino acid transporter heterodimer components SLC7A5 and SLC3A2. These transporters have previously been shown to be essential for TNBC tumorigenesis due to the increased need for glutamine^29,30^. MCF10A (black module) had an enrichment of ribosomal subunits, and MCF-7 (turquoise module) had an enrichment for chromatin and protein-DNA complex proteins. Finally, we observed an enrichment for proteins involved in cell junction and immune response pathways within the MDA-MB-231 (green and blue) modules. Interestingly, p53 was upregulated in MDA-MB-231 when compared to the other three cell lines, consistent with previous proteomic profiling of these cell lines^28,31^, and is vital for MDA-MB-231 survival^31^.

### Relative quantification of KDM abundance in breast cancer cell lines

Next, we compared the quantification of KDMs with and without the KDM trigger channel. Without the trigger channel, only seven KDMs (7/27), or 26% of all KDMs, were quantified (**Fig 2A**). With the addition of the trigger channel, 27 KDMs (100%) were detected, and 26 (96%) were quantified (**Fig 2B**). Seven KDMs were quantified in both experiments (JMJD6, KDM2A, NO66/RIOX1, MINA/RIOX2, KDM1A, KDM3B, KDM5B), and their relative protein abundance was consistent between the two experiments (**Fig 1E**). Three of the seven KDMs quantified in both datasets had a similar number of detected PSMs with or without the trigger channel (KDM1A, KDM3B, and KDM5B). This similarity can be attributed to identical annotated peptide sequences detected in both experiments. However, a close inspection showed that most of the annotated sequences were unique in each experiment. It is also important to note that KDM5B was not detected in the initial GFP-KDM IP emphasizing that stochasticity plays a role in data dependent acquisition mass spectrometry.

**Figure 2:**
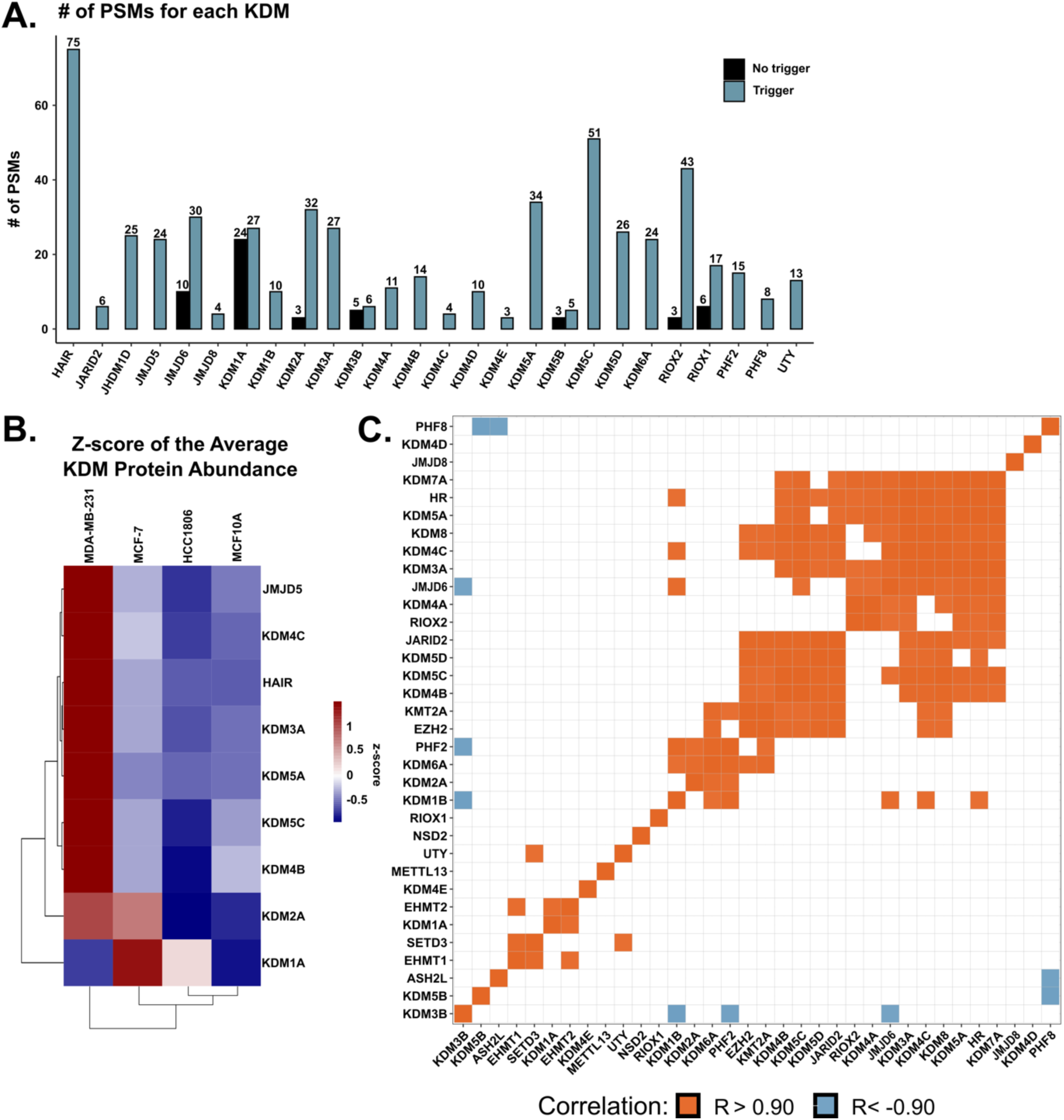
Quantification of lysine demethylases in breast cancer cell lines. **A.** Bar graph depicting the number of PSMs observed with (blue) and without (black) the KDM trigger channel for the indicated KDM. **B.** Heatmap of the z-score of the differentially abundant average protein abundance. Rows are the indicated KDM, while columns are the indicated cell line. Columns and rows are clustered via hierarchical clustering (euclidean distance). **C.** Matrix of the quantified KDMs and KMTs in the trigger channel experiment depicting significant positive (orange) and negative (blue) Pearson correlations of the average protein abundance.

The relative abundance of each KDM within the trigger channel dataset was compared across all four cell lines. Eight KDMs were differentially abundant between the four cell lines (KDM1A, KDM3A, KDM4B, KDM4C, KDM5A, KDM5C, KDM5D, and JMJD5) (ANOVA; *p* < 0.05) (**Fig 2B**). Furthermore, KDM3A, KDM4B, and KDM5C were upregulated in MDA-MB-231, and KDM1A was upregulated in MCF-7 compared to the other three cell lines. The KDM protein abundance in the three cancer cell lines was compared with publicly available mRNA expression (DepMap). Only one positive correlation (RIOX2) was observed between the protein abundance and mRNA expression (**Fig S3**).

Multiple KMTs and KDMs have the same substrate. Therefore, if a cell line has a higher KDM expression, a KMT that methylates the same substrate could be higher to maintain homeostasis. This balance may not occur for all KMTs and KDMs for various reasons, such as disease-related abnormal protein expression. Eight KMTs were quantified and were included in the analysis to make connections between KMT and KDM expression. The expression of several KDMs significantly correlated with one another (**Fig 2C**, R> |0.9|; *p* < 0.05). For example, KDM1B negatively correlated with KDM3B and positively correlated with KDM6A, PHF2, JMJD6, KDM4C, and HR. We also observed a positive correlation between some KMTs and KDMs, including EHMT2 and KDM1A. Interestingly, EHMT2 methylates H3K9me1/2 while KDM1A demethylates the same site, suggesting similar expression levels between these two proteins due to their activity on a shared substrate. In addition, EZH2 positively correlated with two KDMs (KDM6A and JARID2) that demethylate H3K27me3, the product of EZH2. This is expected, as JARID2 and EZH2 are components of the polycomb repressive complex 2 (PRC2). The observed correlations between KMT and KDM expression showcase the potential utility of the trigger channel to discover and validate KMT and KDM signaling networks.

### TMT labeling enables the quantification of hundreds of lysine methylation sites within a single mass spectrometry experiment

After the characterization of protein abundances, we focused on detecting and quantifying Kme sites. Previous work demonstrated that antibody enrichment is unnecessary for detecting Kme sites. This study investigated whether detection and quantification of Kme sites within a TMT-based proteomics workflow was feasible. For this analysis, we looked at all Kme sites identified and quantified in both experiments (with and without the KDM trigger channel). In total, 608 unique Kme sites on 387 unique proteins were detected, with 578 being novel (PhosphoSitePlus©)^32^ (**Fig 3A**). From the 608 detected Kme sites, 326 unique Kme peptides were quantified from 203 unique proteins, of which 303 were novel (**Fig 3B**).

**Figure 3:**
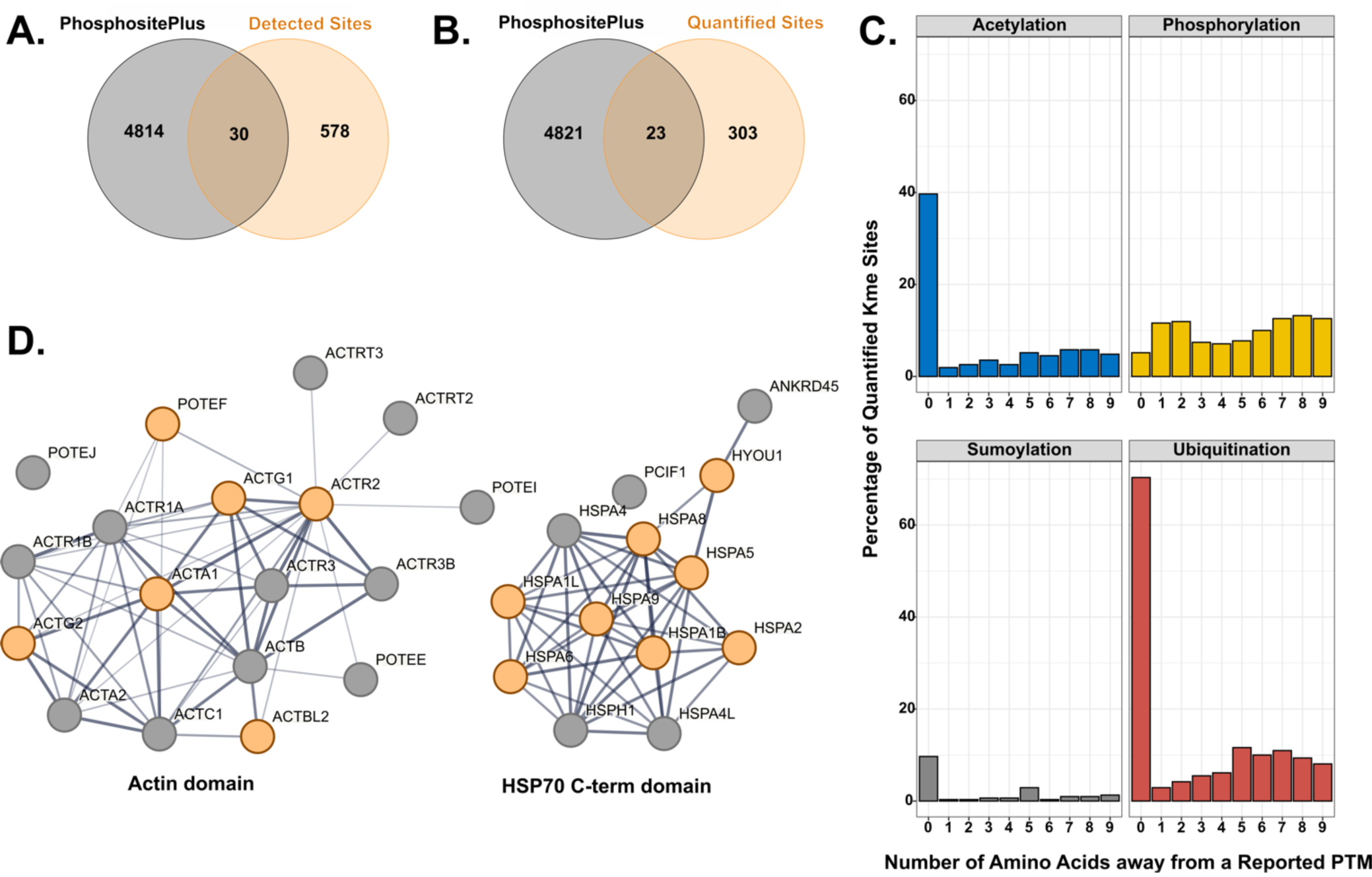
Characterization of the quantified lysine methylation sites. Venn diagram of previously observed Kme sites from PhosphoSitePlus© and those detected (**A**) or quantified (**B**) within this study. **C.** Percent of quantified Kme sites that either share the same site (position = 0) with or are within nine amino acids (position = 1-9) from observed acetylation, ubiquitination, sumoylation, or phosphorylation sites as annotated in PhosphoSitePlus© **D.** STRING network diagrams of domains enriched for lysine methylation. Orange circles represent proteins with a quantified Kme site within this study.

Next, we compared how the addition of the isobaric KDM trigger channel impacted the detection or quantification of Kme sites. More sites were detected and quantified without the inclusion of the trigger channel, though a substantial number of sites were still identified with the addition of the trigger channel (408 vs. 340 identified; 239 vs. 173 Kme sites quantified). A total of 53 Kme sites were quantified in both datasets. A Pearson correlation analysis between the two experiments revealed adequate correlation (R > 0.4; *p* < 0.05), suggesting reproducible Kme site quantification (**Fig S4A**). Overall, these data suggest that the isobaric KDM trigger channel did not significantly impact the detection and quantitation of lysine methylation sites.

In addition to methylation, lysine can be modified by other PTMs, including acetylation, sumoylation, or ubiquitination. Other studies have noted the antagonistic or symbiotic relationship, depending on the cellular context, between lysine and phosphorylation ^33,34^. We, therefore, queried how many Kme sites had reported phosphorylation, sumoylation, ubiquitination, or acetylation site either on that same lysine or within nine amino acids of the methylated lysine (PhosphoSitePlus©) (**Fig 3C**). In our study, 40% of the lysines with a quantified Kme site also have been observed to be acetylated, while 10% have been observed to be sumoylated. Interestingly, 70% can also be ubiquitinated, suggesting a strong potential connection between methylation and ubiquitination. Phosphorylation has an equal distribution across all nine sites, with an average of 10% of Kme sites with a neighboring phosphorylation site.

Next, we performed a Gene Ontology enrichment analysis on all the lysine methylated proteins. There was an enrichment for proteins involved in protein folding, cytoskeletal organization, and junctions, all of which are terms that have been previously associated with lysine methylation (**Fig S4B**). Interestingly, two ATP binding domains had an enrichment for methylated proteins: actin (6/18) and HSP70 C-term (8/13) domains (**Fig 3D**). While most of these Kme sites are novel, their roles are consistent with previously profiled Kme sites observed in different cell types. These findings emphasize the integral and diverse role of lysine methylation.

### Breast cancer cell lines have unique non-histone lysine methylome signatures

Histone lysine methylation sites have unique profiles across different breast cancer subtypes^35^. We investigated if the same was true for non-histone lysine methylation sites. To identify differentially abundant Kme sites, the Kme peptide abundance was normalized to the protein abundance. Of the 326 unique Kme sites, 142 were significantly differentially abundant between all four cell lines (ANOVA, *p* > 0.05) (**Fig 4**). WGCNA was conducted on the differentially abundant sites. Three distinct clusters were identified: blue, brown, and turquoise (**Fig. S5 A & B**). Proteins involved in protein folding, secretion, and cytoskeletal structure were enriched in MDA-MB-231 (turquoise and brown), while cytoskeletal and cell junction proteins were enriched in MCF7 (blue) (GO term analysis) (**Fig 4**.). Therefore, similar to histone lysine methylation sites, non-histone sites have unique and cell type-specific profiles.

**Figure 4:**
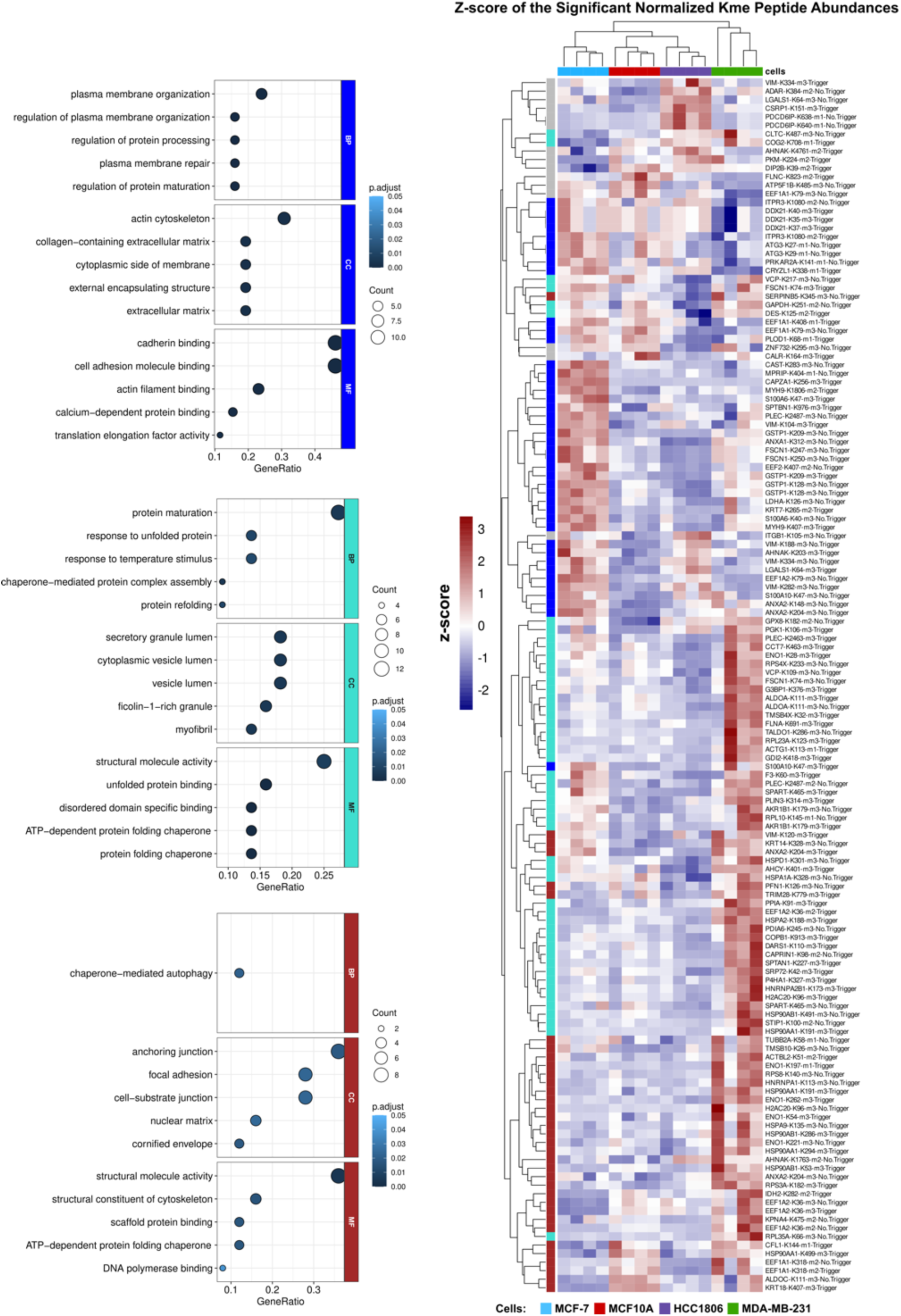
Lysine methylation has distinct profiles across breast cancer cell lines. Heatmap of the differentially expressed Kme sites (*n* =142). Colors represent the z-score of the normalized Kme peptide abundance. Rows (Kme sites) and columns (cell line) are clustered by euclidian distance. Module colors indicate the module to which the Kme site belongs. Enriched GO terms found in the indicated modules are depicted on the left (blue, turquoise, brown) (*p*<0.05).

Overall, 52 Kme sites were significantly upregulated or downregulated within a particular cell line (**Fig S6**). MCF-7 and MDA-MB-231 cells contained the largest number of upregulated sites. A few sites were downregulated in MCF10A, but no single site was more abundant in MCF10A compared to the three other cell lines. Interestingly, two sites were upregulated in all the breast cancer cell lines: ADAR K384-me2 and GPX8 K182-me2 (**Fig 5A & B**). Double-stranded RNA-specific adenosine deaminase (ADAR) is a deaminase that catalyzes the hydrolytic deamination of adenosine to inosine, which has been shown to impact overall RNA regulation and downstream protein expression. Intriguingly, there have been studies connecting the overexpression of ADAR to breast cancer progression^36^. Knockdown of ADAR led to a decrease in triple-negative breast cancer cell proliferation and overall transformation and tumorigenesis in TNBC xenograft models^32^. GPX8 (Probable glutathione peroxidase 8) has been connected to breast cancer metastasis, with a recent study showing MDA-MB-231 cells lacking GPX8 were less invasive and reverted from a mesenchymal to an epithelial state^37^.

**Figure 5:**
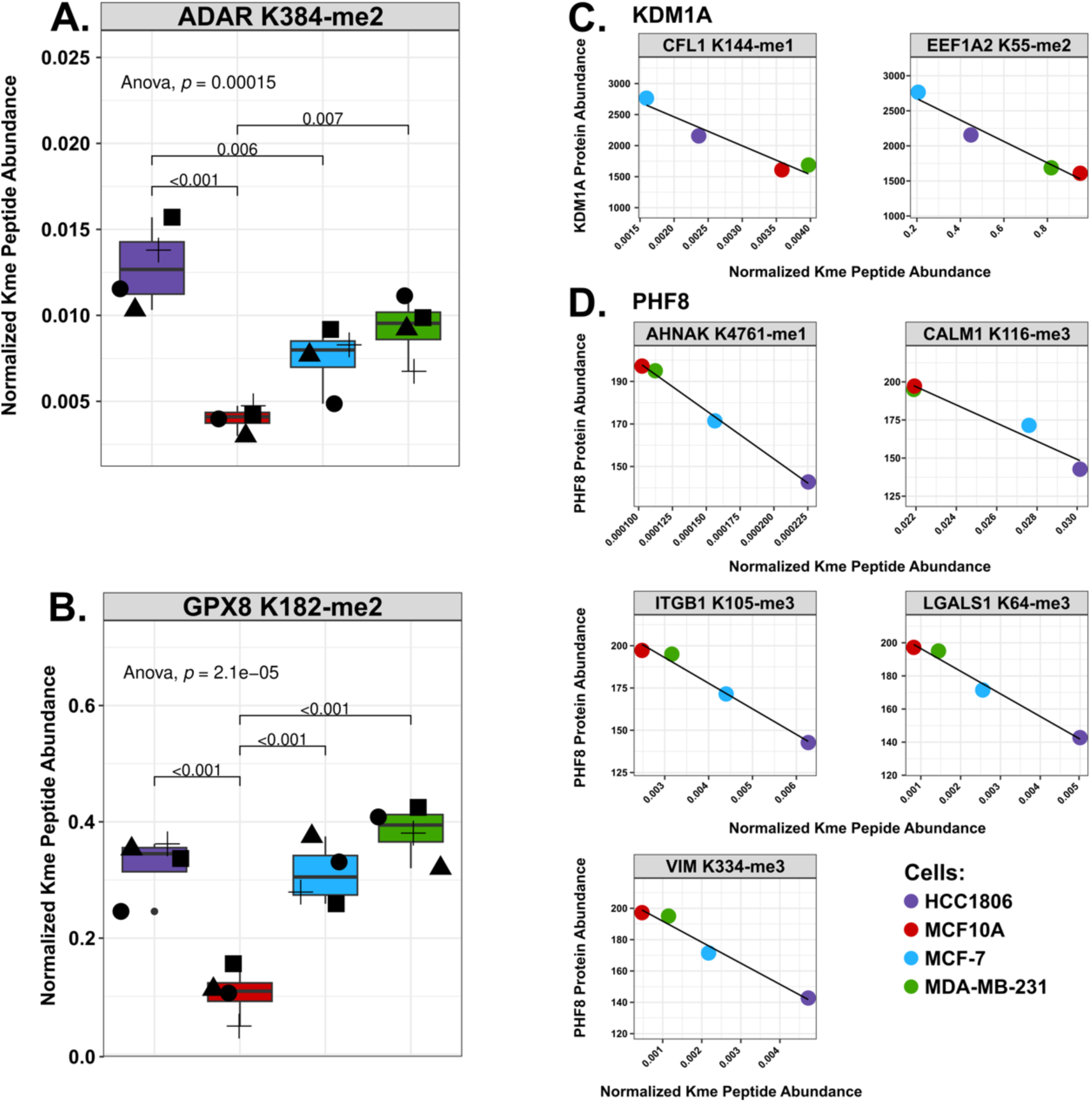
Comparison of normalized Kme site abundances and correlation with KDM levels. **A.** Boxplots of the normalized Kme site abundances (y-axis) of ADAR K384-me2 and GPX8 K128-me2 in HCC1806 (purple), MCF10A (red), MCF-7 (red), and MDA-MB-231 (green). Shapes represent different biological replicates. **B.** Correlation between the protein abundance for indicated KDMs (KDM1A or PHF8) and normalized Kme peptide abundance as indicated (gray box).

To explore potential regulatory mechanisms, we investigated whether any KDMs correlated with specific Kme sites. Since KDMs demethylate Kme sites, we reasoned that focus should be placed on sites that negatively correlated with a KDM, as a greater abundance of the KDM would lead to a lower abundance of a methylated substrate. All 26 KDMs negatively correlated with at least one Kme site (**Fig S7**). For example, KDM1A (LSD1) negatively correlated with CFL1 K144 (Cofilin-1) and EEF1A2 K55 (Elongation factor 1-alpha 2), with higher abundance in MCF-7 and lower abundance in MDA-MB-231 or MCF10A (**Fig 5C**). EEF1A1/2 K55-me2 is catalyzed by the KMT METTl13 and is associated with tumor proliferation in lung and pancreatic cancer cells^3,38^. PHF8 negatively correlated with AHNAK K4761 me1 (Neuroblast differentiation-associated protein AHNAK), CALM1 K116 me3 (Calmodulin-1), ITGB1 K105 (Integrin beta-1), LGALS1 K64 (Galectin-1), and VIM K334 me3 (Vimentin), which all have a higher expression within MCF10A and MDA-MB-231 and the lowest expression in HCC1806 (**Fig 5D**). However, the motifs of the negatively correlated sites did not have any striking similarities (**Table S2**). Future work capturing the expression of KMTs, KDMs, and Kme sites in additional cell lines and tissues will help create a systems-level understanding of lysine methylation signaling networks and facilitate the identification of putative enzyme-substrate connections.

## DISCUSSION

In this work, we simultaneously profile the abundance of lysine demethylases and lysine methylation sites utilizing Tandem Mass Tag labels and a novel KDM isobaric trigger channel. Previous attempts to connect methyl regulators and Kme sites relied on SILAC and antibody enrichment. While successful, this required an appreciable amount of material per enrichment (20 milligrams), expensive enrichment reagents, and an individual MS run for each enriched fraction, making it more costly and less reproducible. Furthermore, the addition of an isobaric trigger channel to quantify KDMs simultaneously is not feasible in such a workflow. To ensure that methyl mediators are reproducibly detected and quantified, we generated a KDM isobaric trigger channel. Others have demonstrated the effective use of trigger channels to boost the detection of low-abundance proteins using MS^19–21^. Indeed, using the KDM trigger channel enabled the quantification of 96% (26 of 27) of all human KDMs compared to 26% (7 of 27) without the trigger channel. The use of the trigger channel revealed a higher abundance of KDM3A, KDM4B, and KDM5C within MDA-MB-231 and KDM1A in MCF7. Previous studies have shown these KDMs are integral to breast cancer tumorigenesis^39–44^. Furthermore, one group demonstrated that KDM3A has one non-histone target in the MDA-MB-231 cell line: p53 K372 me1^39^. While we did not detect p53 K372 within this study, this workflow can be used to expand our current understanding of KDMs and their non-histone substrates.

Previous work from our lab and others has demonstrated successful detection of Kme sites without antibody enrichment^1,45^. Using TMT labeling, we demonstrate that Kme sites can be quantitatively profiled sans antibody enrichment within a single MS run. When combining both MS experiments (with and without the trigger channel), 326 total Kme sites were quantified. Over 50 of the same Kme sites were quantified in both experiments. While this constitutes only a 30% overlap between the two runs, previous work using antibody enrichment observed only a 20% overlap between enrichment strategies^1^. With the ability to quantify Kme sites across multiple samples within a single MS run, we reveal that, similar to histone Kme sites, breast cancer cell lines have unique non-histone Kme site profiles. The quantified Kme sites are found on proteins involved in processes previously connected to regulation by lysine methylation, including scaffolding proteins and molecules involved in protein folding. Interestingly, there is an enrichment for Kme sites involved in distinct processes in the individual cell lines. In conjunction with the observed KDM profile, this suggests cell-type specific lysine methylation signaling networks.

Given the ability to profile both KDMs and Kme sites across four lines, we also show that the abundance of different KDMs negatively correlates with different Kme sites. While more work needs to be done to delineate whether these are putative substrates, using this platform in various cell lines and with various conditions, including drug treatments, can aid in connecting lysine methylation regulators and their putative non-histone substrates. Furthermore, this workflow can be applied to observe Kme site abundance changes and potential compensatory mechanisms upon genetic manipulation or inhibition of specific lysine methylation mediators.

## Data Availability

Code used to analyze this data is accessible at https://github.com/caberryhill/TNBC-Analysis. All raw mass spectrometry data and processed Proteome Discoverer result files are uploaded to MassIVE repository with accession MSV000095132.

## Acknowledgements

The authors thank members of the Cornett and Mosley labs for helpful feedback and discussion. The mass spectrometry work was performed by the Indiana University School of Medicine (IUSM) Center for Proteome Analysis (CPA). Acquisition of the IUSM CPA instrumentation used for this project was provided in part by the Indiana University Precision Health Initiative and the IU Simon Comprehensive Cancer Center. The proteomics work was supported, in part, by the Indiana Clinical and Translational Sciences Institute (Award Number UL1TR002529 from the National Institutes of Health, National Center for Advancing Translational Sciences, Clinical and Translational Sciences Award) and, in part, by the IU Simon Comprehensive Cancer Center Support Grant (Award Number P30CA082709 from the National Cancer Institute). The research reported in this publication was supported by the National Institute of General Medical Sciences of the National Institutes of Health under award number R35GM147023 (EMC). The content is solely the responsibility of the authors and does not necessarily represent the official views of the National Institutes of Health.

## Author Contributions

The manuscript was written with contributions from all authors. All authors have given approval to the final version of the manuscript.

