## Supporting Info for "Quantitative analysis of non-histone lysine methylation sites and lysine demethylases in breast cancer cell lines"

**The PDF file includes:**

Figure S1. Reproducible quantification of the breast cancer proteomes.
Figure S2. Differential protein abundances reveal differences between the breast cell lines.
Figure S3: Few significant correlations between KDM mRNA expression and protein abundances.
Figure S4: Addition of the trigger channel does not impact Kme site quantification.
Figure S5: WGCNA analysis reveals distinct clusters.
Figure S6: Kme sites upregulated or downregulated in a cell-specific manner.
Figure S7: KDM and Kme site correlations.
Supporting Table 1: Summary of LC-MS/MS data with and without the isobaric trigger channel.
Supporting Table 2: Motifs of negatively correlated Kme sites.

**Other supporting info includes:**

Supporting Table 3 (excel file): Tables with quantified proteins and sites.

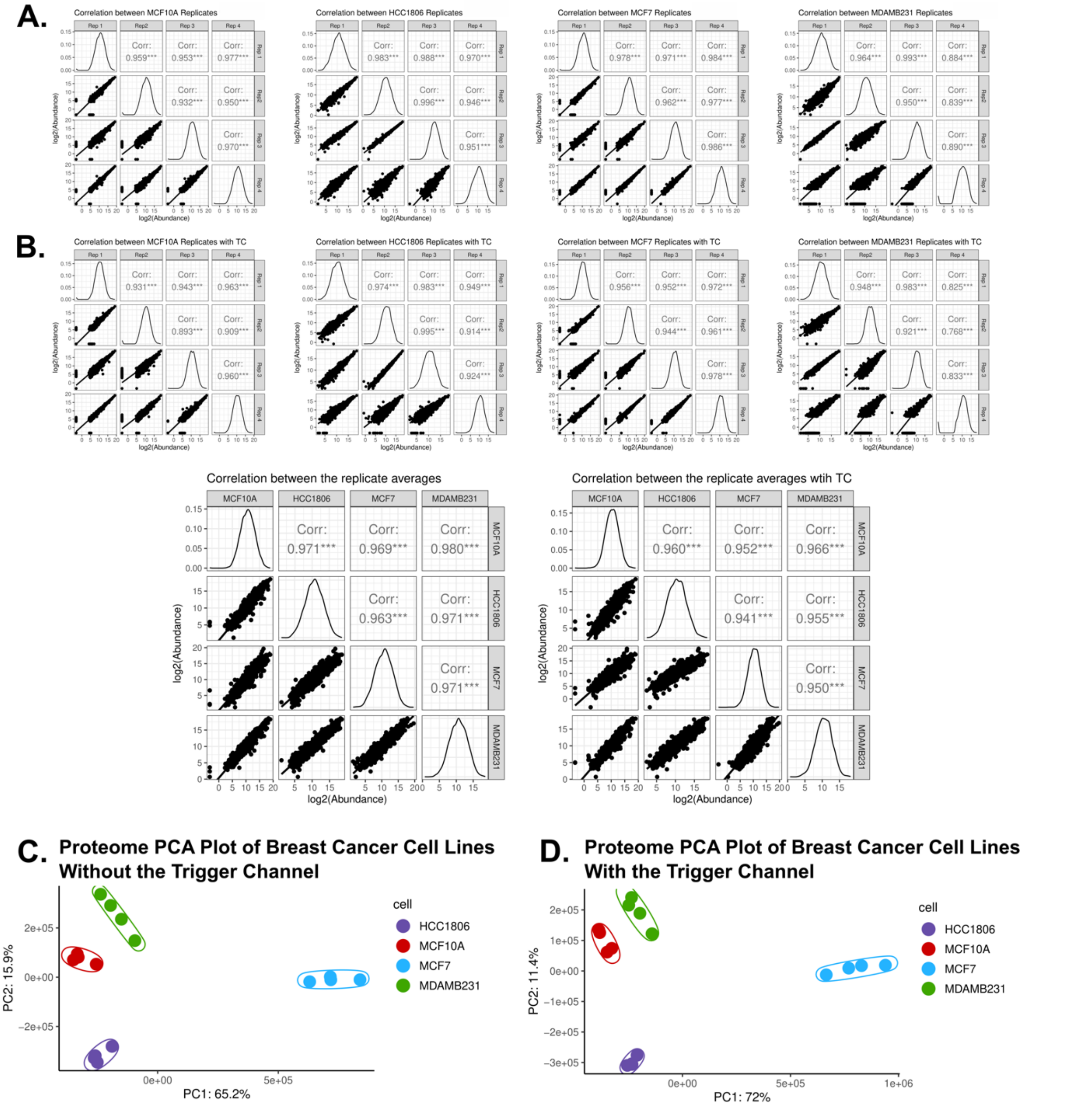


**Figure S1. Reproducible quantification of the breast cancer proteomes.**
Pearson correlation plots of the biological replicates within each experiment **A.** without the trigger channel and **B.** with the trigger channel (***= $p < 0.001$). Principal component analysis (PCA) of the breast cancer proteomes **C.** without or **D.** with the trigger channel. Each colored dot is a biological replicate of the indicated cell line.

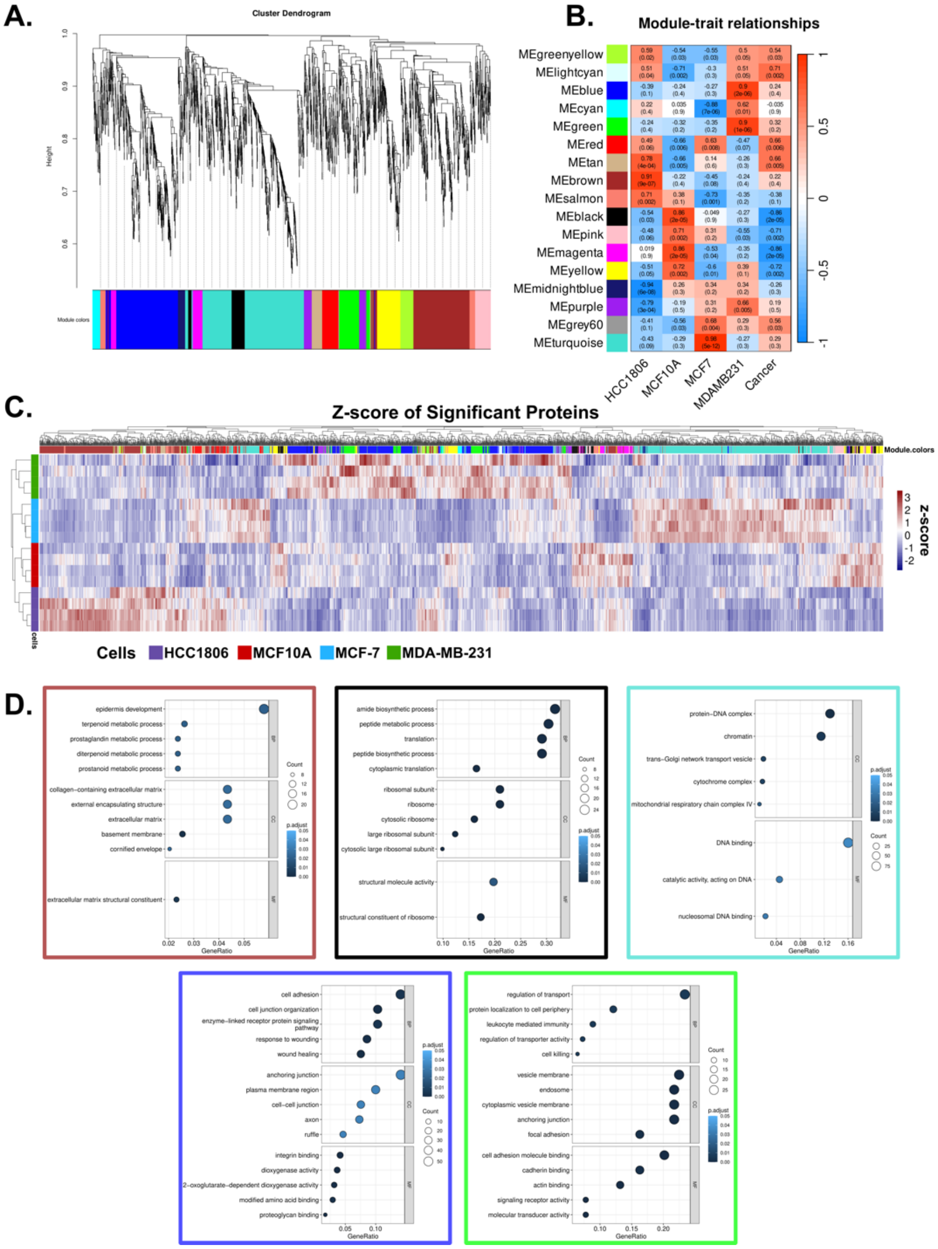

**Figure S2. Differential protein abundances reveal differences between the breast cell lines.**
**A.** WGCNA dendrogram and **B.** identified modules from the differentially abundant proteins from the trigger channel experiment **C.** Heatmap of the differentially abundant proteins ($n$ = 2,290; ANOVA, $p < 0.05$). Colors represent the z-score of the protein abundances. Euclidean distance was used to cluster the rows (cell replicates) and columns (proteins). The module color associated with a given protein is represented along the top. **D.** Enriched GO terms within the top significant cluster corresponding to a particular cell line ($p < 0.05$). The brown module corresponds to HCC1806, blue and green correspond to MDA-MB-231, black is MCF10A, and turquoise is MCF-7.

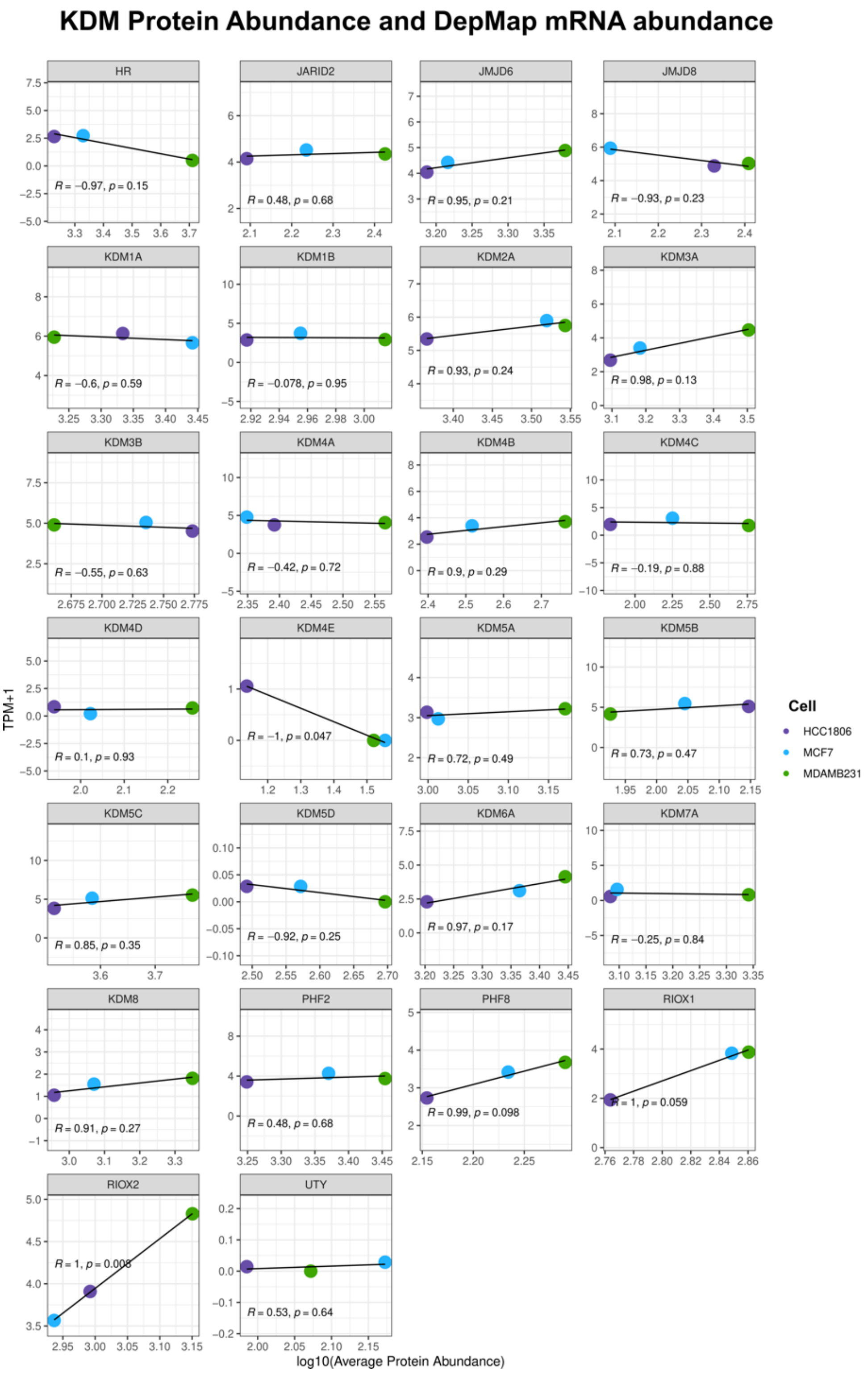


**Figure S3: Few significant correlations between KDM mRNA expression and protein abundances.**

Scatter plots of the $\log_{10}$ KDM average protein expression observed in the trigger channel (x-axis) and DepMap mRNA expression (y-axis). The colored points correspond to the indicated cell line, and the KDM protein name is found in the gray box above the graph. Pearson correlation and *p*-values are listed within the plot.

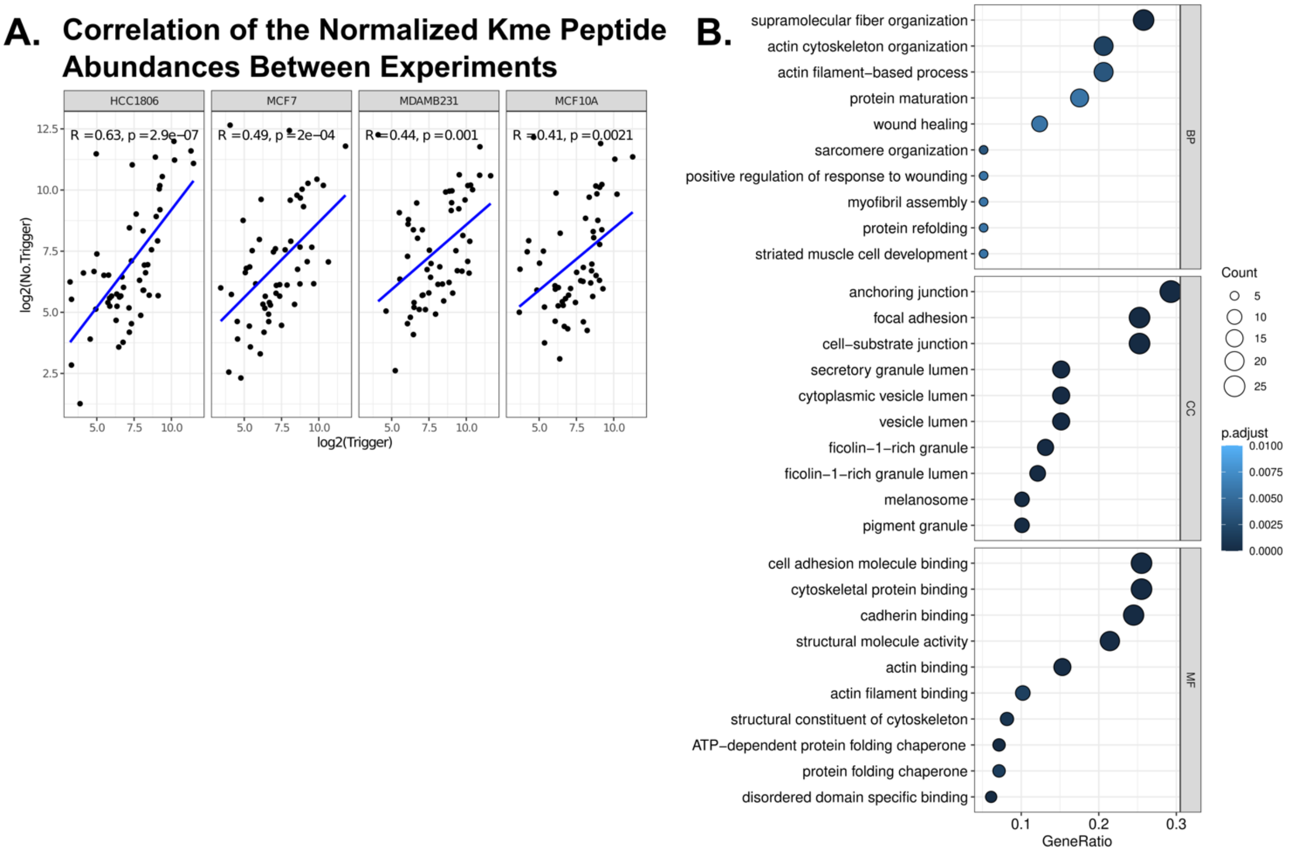


**Figure S4: Addition of the trigger channel does not impact Kme site quantification.** **A.** Pearson correlation of the normalized Kme peptide abundances with (x-axis) and without (y-axis) the trigger channel ($n$ = 53) divided by the indicated cell line. **B.** Enriched GO terms (biological processes, molecular function, and cellular compartments) of the quantified lysine methylated proteins (adjusted $p$-value < 0.05).

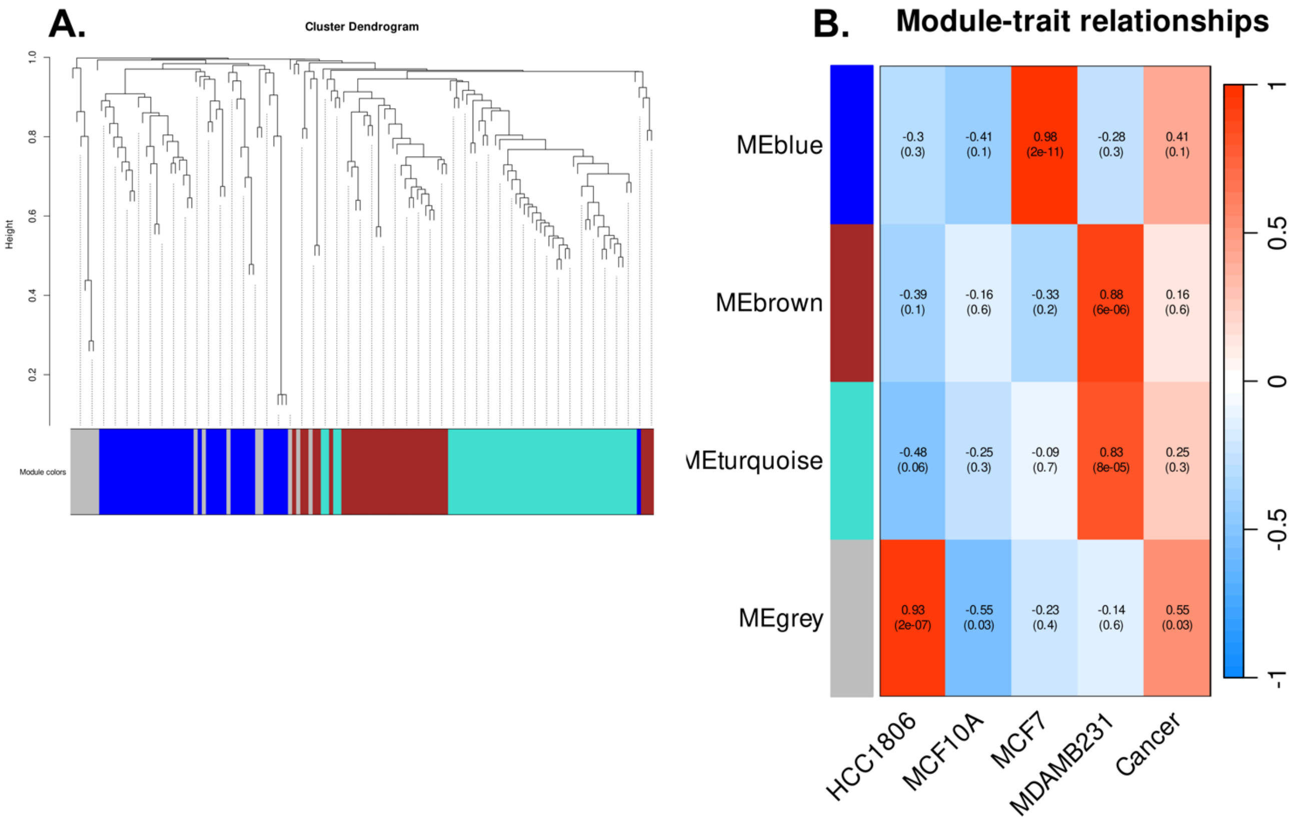


**Figure S5: WGCNA analysis reveals distinct clusters.**
**A.** WGCNA dendrogram and identified modules from the differentially abundant Kme peptides (ANOVA; $p < 0.05$). Significant modules have $p < 0.05$.

### Berryhill, C. et al. 2024. Fig. S6

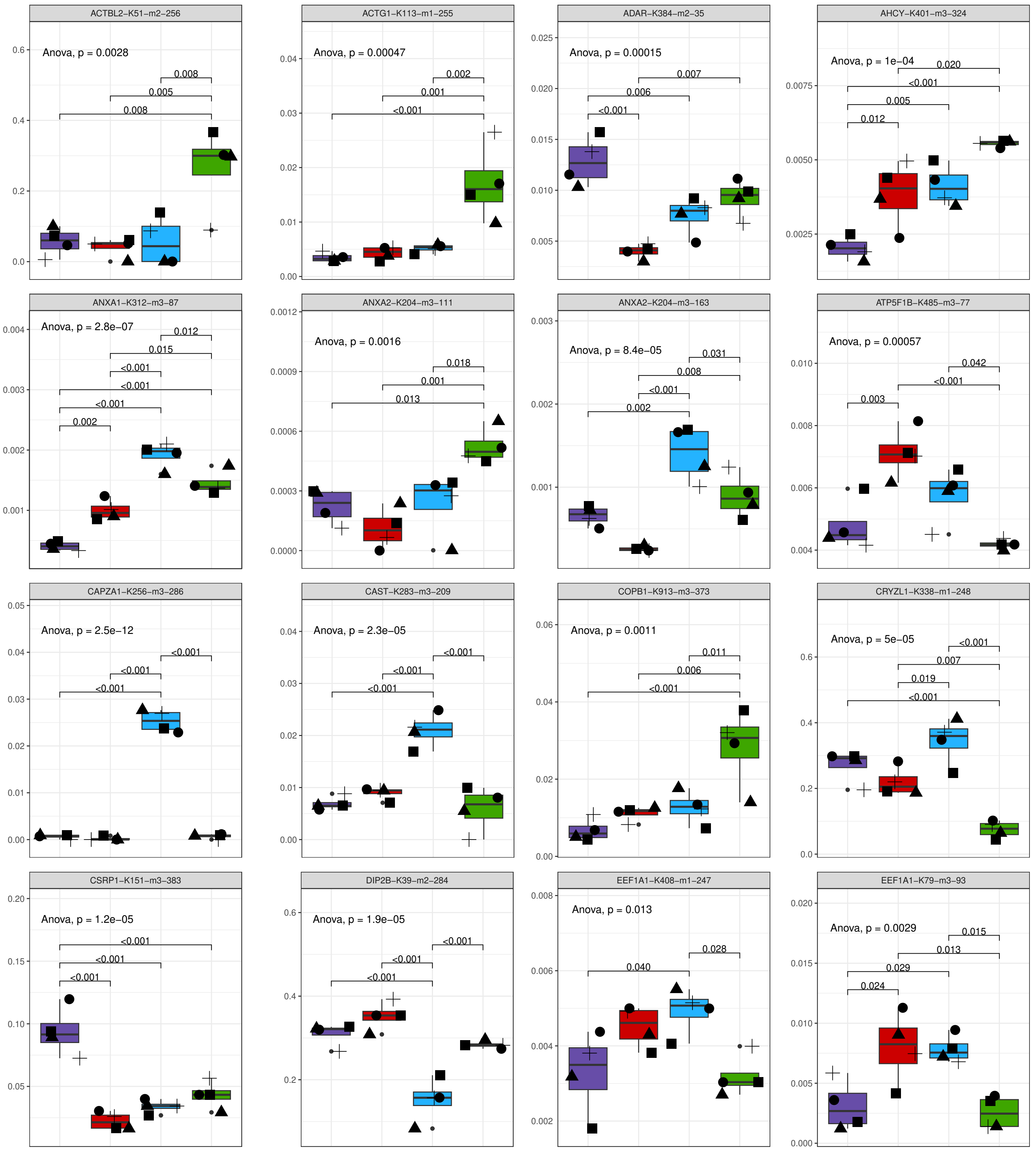

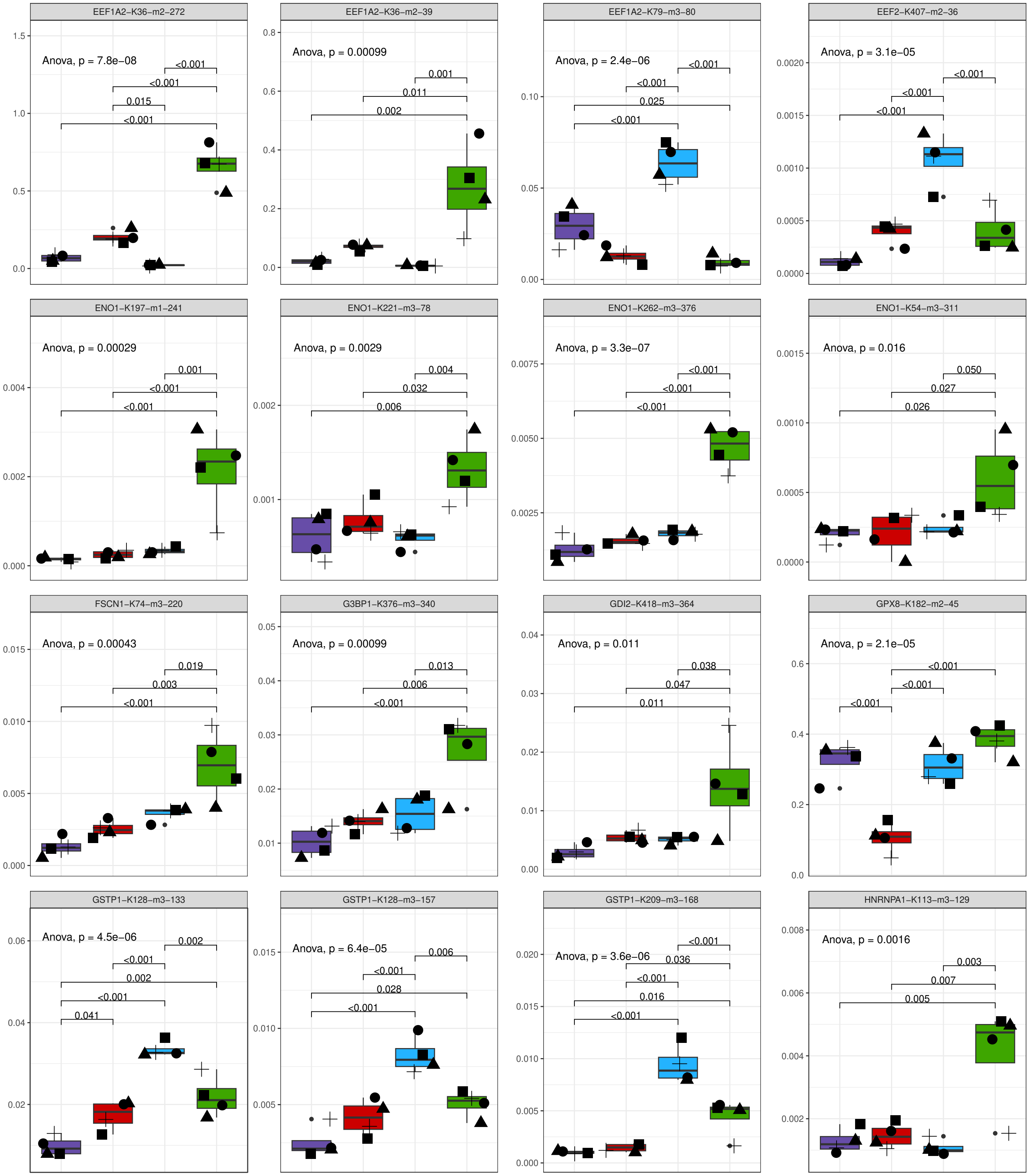

EEF1A2–K36–m2–272
Anova, p = 7.8e−08
<0.001
<0.001
0.015
<0.001
EEF1A2–K36–m2–39
Anova, p = 0.00099
0.001
0.011
0.002
EEF1A2–K79–m3–80
Anova, p = 2.4e−06
<0.001
<0.001
0.025
<0.001
EEF2–K407–m2–36
Anova, p = 3.1e−05
<0.001
<0.001
<0.001
ENO1–K197–m1–241
Anova, p = 0.00029
0.001
<0.001
<0.001
ENO1–K221–m3–78
Anova, p = 0.0029
0.004
0.032
0.006
ENO1–K262–m3–376
Anova, p = 3.3e−07
<0.001
<0.001
<0.001
ENO1–K54–m3–311
Anova, p = 0.016
0.050
0.027
0.026
FSCN1–K74–m3–220
Anova, p = 0.00043
0.019
0.003
<0.001
G3BP1–K376–m3–340
Anova, p = 0.00099
0.013
0.006
<0.001
GDI2–K418–m3–364
Anova, p = 0.011
0.038
0.047
0.011
GPX8–K182–m2–45
Anova, p = 2.1e−05
<0.001
<0.001
<0.001
GSTP1–K128–m3–133
Anova, p = 4.5e−06
0.002
<0.001
0.002
<0.001
0.041
GSTP1–K128–m3–157
Anova, p = 6.4e−05
0.006
<0.001
0.028
<0.001
GSTP1–K209–m3–168
Anova, p = 3.6e−06
<0.001
0.036
<0.001
0.016
<0.001
HNRNPA1–K113–m3–129
Anova, p = 0.0016
0.003
0.007
0.005

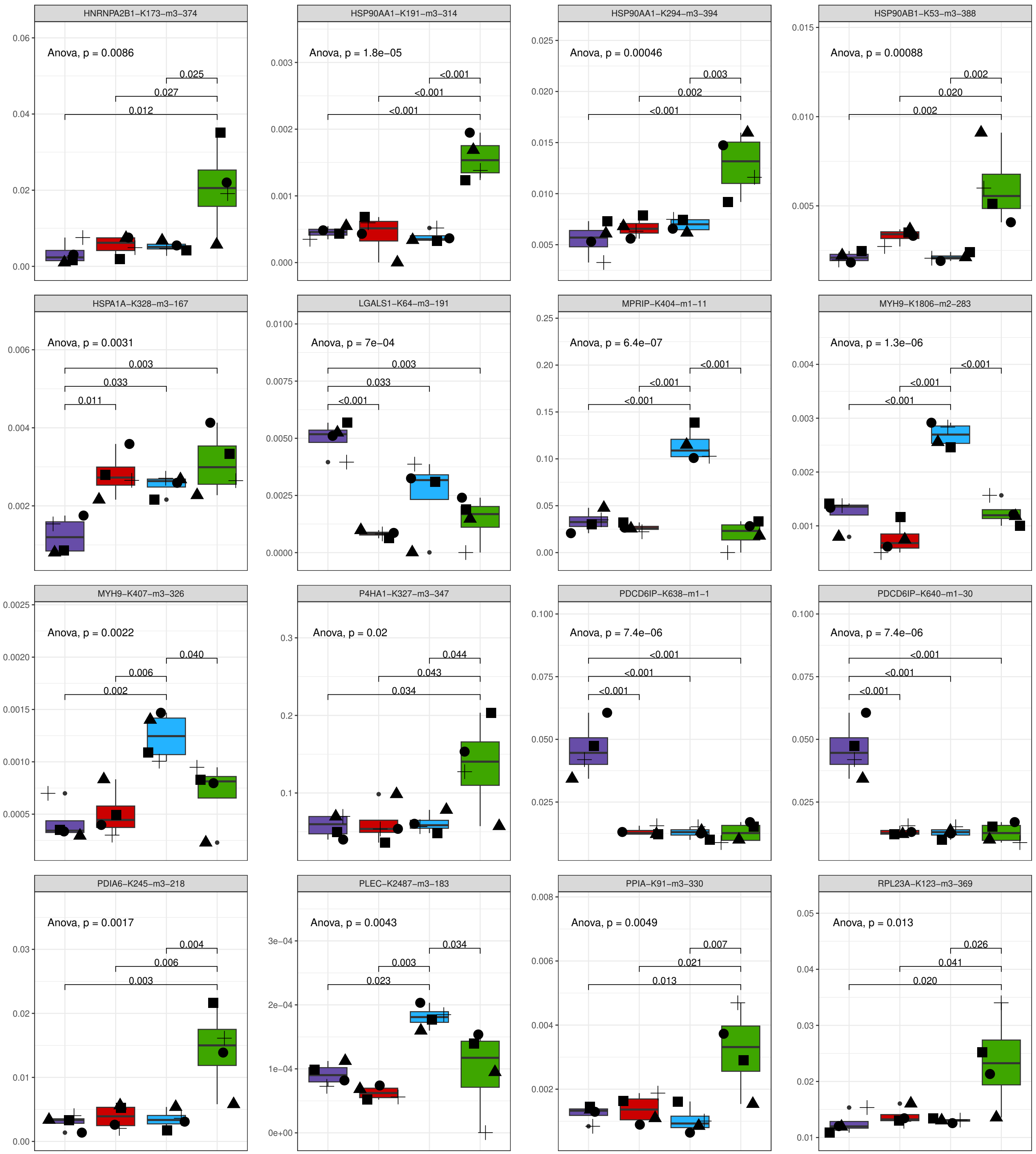
HNRNPA2B1−K173−m3−374
Anova, p = 0.0086
0.025
0.027
0.012
HSP90AA1−K191−m3−314
Anova, p = 1.8e−05
<0.001
<0.001
<0.001
HSP90AA1−K294−m3−394
Anova, p = 0.00046
0.003
0.002
<0.001
HSP90AB1−K53−m3−388
Anova, p = 0.00088
0.002
0.020
0.002
HSPA1A−K328−m3−167
Anova, p = 0.0031
0.003
0.033
0.011
LGALS1−K64−m3−191
Anova, p = 7e−04
0.003
0.033
<0.001
MPRIP−K404−m1−11
Anova, p = 6.4e−07
<0.001
<0.001
<0.001
MYH9−K1806−m2−283
Anova, p = 1.3e−06
<0.001
<0.001
<0.001
MYH9−K407−m3−326
Anova, p = 0.0022
0.040
0.006
0.002
P4HA1−K327−m3−347
Anova, p = 0.02
0.044
0.043
0.034
PDCD6IP−K638−m1−1
Anova, p = 7.4e−06
<0.001
<0.001
<0.001
PDCD6IP−K640−m1−30
Anova, p = 7.4e−06
<0.001
<0.001
<0.001
PDIA6−K245−m3−218
Anova, p = 0.0017
0.004
0.006
0.003
PLEC−K2487−m3−183
Anova, p = 0.0043
0.034
0.003
0.023
PPIA−K91−m3−330
Anova, p = 0.0049
0.007
0.021
0.013
RPL23A−K123−m3−369
Anova, p = 0.013
0.026
0.041
0.020

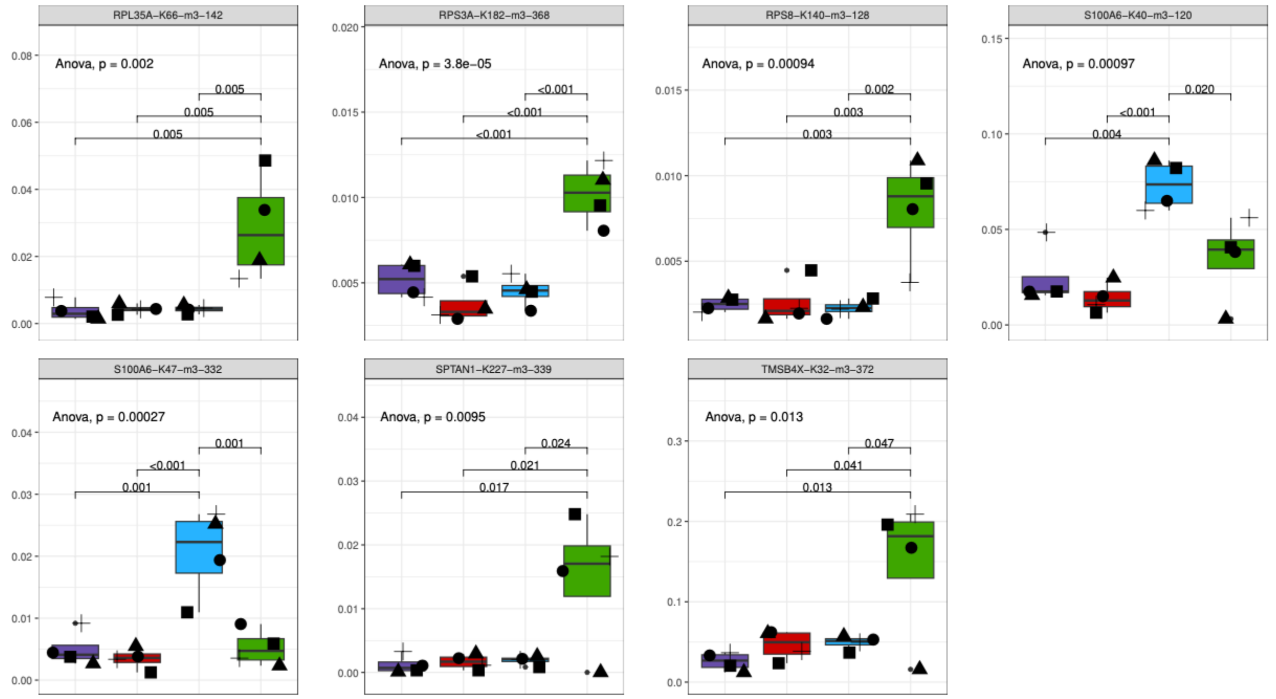


**Figure S6: Kme sites upregulated or downregulated in a cell-specific manner.** Boxplots of the normalized Kme site abundances (y-axis) ($n$ = 52). Purple indicates HCC1806, red is MCF10A, blue is MCF-7, and green is MD-AMB-231.

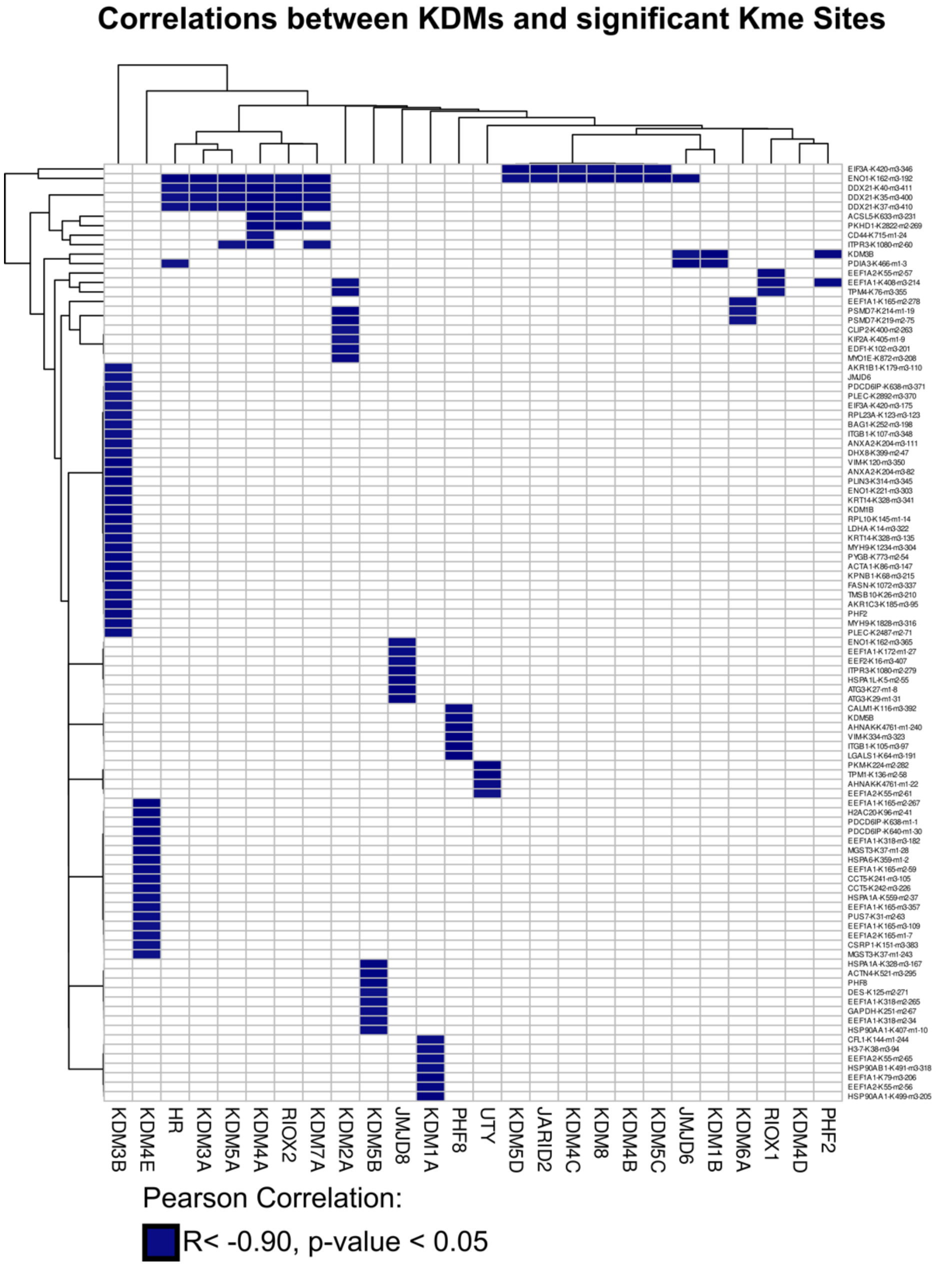


**Figure S7: KDM and Kme site correlations.**
Heatmap depicting the Pearson correlation values between average KDM protein abundance and the normalized Kme site abundance. Only significant and negative correlations are visualized. Columns are KDMs and rows are Kme sites.

**Supporting Table 1: Summary of LC-MS/MS data with and without the isobaric trigger channel.**

| | Without Trigger | With Trigger |
|---|---|---|
| # of PSMs | 142,475 | 126,891 |
| # of Peptides | 54,727 | 47,699 |
| # of proteins | 4,200 | 4,384 |
| # of quantified proteins | 4,131 | 4,311 |

**Supporting Table 2: Motifs of negatively correlated Kme sites.**

| | Site | Motif |
|---|---|---|
| KDM1A | CLF K144 | EEVKDRC |
| | EEF1A2 K55 | GSFKYAW |
| PHF8 | AHNAK K4761 | KGPKVDI |
| | ITGB1 K105 | TAEKLKP |
| | LGALS1 K64 | CNSKDGG |
| | VIM K334 | DALKGTN |
| | CALM1 K116 | LGEKLTD |
